# Transcriptome analysis of hippocampal subfields identifies gene expression profiles associated with long-term active place avoidance memory

**DOI:** 10.1101/2020.02.05.935759

**Authors:** Rayna M. Harris, Hsin-Yi Kao, Juan Marcos Alarcón, André A. Fenton, Hans A. Hofmann

## Abstract

The hippocampus plays a critical role in storing and retrieving spatial information. By targeting the dorsal hippocampus and manipulating specific “candidate” molecules using pharmacological and genetic manipulations, we have previously discovered that long-term active place avoidance memory requires transient activation of particular molecules in dorsal hippocampus. These molecules include amongst others, the persistent kinases Ca-calmodulin kinase II (CaMKII) and the atypical protein kinase C isoform PKC *ι/λ* for acquisition of the conditioned behavior, whereas persistent activation of the other atypical PKC, protein kinase M zeta (PKM*ζ*) is necessary for maintaining the memory for at least a month. It nonetheless remains unclear what other molecules and their interactions maintain active place avoidance long-term memory, and the candidate molecule approach is both impractical and inadequate to identify new candidates since there are so many to survey. Here we use a complementary approach to identify candidates by transcriptional profiling of hippocampus subregions after formation of the long-term active place avoidance memory. Interestingly, 24-h after conditioning and soon after expressing memory retention, immediate early genes were upregulated in the dentate gyrus but not Ammon’s horn of the memory expressing group. In addition to determining what genes are differentially regulated during memory maintenance, we performed an integrative, unbiased survey of the genes with expression levels that covary with behavioral measures of active place avoidance memory persistence. Gene Ontology analysis of the most differentially expressed genes shows that active place avoidance memory is associated with activation of transcription and synaptic differentiation in dentate gyrus but not CA3 or CA1, whereas hypothesis-driven candidate molecule analyses identified insignificant changes in the expression of many LTP-associated molecules in the various hippocampal subfields, nor did they covary with active place avoidance memory expression, ruling out strong transcriptional regulation but not translational regulation, which was not investigated. These findings and the data set establish an unbiased resource to screen for molecules and evaluate hypotheses for the molecular components of a hippocampus-dependent, long-term active place avoidance memory.

## Introduction

The hippocampus is a thoroughly studied model system for investigating how spatial memory is stored, and it has been a valuable platform for candidate molecule approaches to investigating the cellular mechanisms of memory formation and persistence. For example, by targeting and manipulating specific candidate molecules using pharmacological and genetic manipulations, it was discovered that active place avoidance memory requires transient activation of several long-term potentiation (LTP)-associated molecules in dorsal hippocampus, including a number of kinases such as the persistent kinases Ca-calmodulin kinase II (CaMKII) and the atypical protein kinase C isoform PKC/*ι/λ* for acquisition of the conditioned behavior (Tsokas *et al.*, 2016; Rossetti *et al.*, 2017; Sacktor and Fenton, 2018). Importantly, these molecules are not critical for maintaining the memory, whereas persistent activation of the other atypical PKC, protein kinase M zeta (PKM*ζ*) lasting at least a month is necessary for long-term memory (Pastalkova *et al.*, 2006; Tsokas *et al.*, 2016; Hsieh *et al.*, 2017), as shown in other memory systems (Drier *et al.*, 2002; Shema, Sacktor and Dudai, 2007; Serrano *et al.*, 2008; von Kraus, Sacktor and Francis, 2010; Hardt *et al.*, 2010; Madroñal *et al.*, 2010; Migues *et al.*, 2010; Cai *et al.*, 2011; LaFerriere *et al.*, 2011; Shema *et al.*, 2011; Barry *et al.*, 2012; Vélez-Hernández *et al.*, 2013; Li *et al.*, 2014; Wang *et al.*, 2016).

Training in the active place avoidance task changes hippocampal synaptic function as well as the dynamics of local field potentials and the activity of place cells in the hippocampus (Bures *et al.*, 1997; Cimadevilla *et al.*, 2000; Pavlowsky *et al.*, 2017). The active place avoidance memory is maintained by persistent PKM*ζ* activity (Hsieh *et al.*, 2017). Memory-mediating molecules, such as PKM*ζ* that are crucial for memory maintenance, interact with other effector and regulatory molecules in ways that remain largely unknown (Kelly *et al.*, 2007; Kelly, Crary and Sacktor, 2007; Yao *et al.*, 2008; Tian *et al.*, 2008; Migues *et al.*, 2010; Sacktor, 2011; Kandel, 2012; Shao *et al.*, 2012; Eom *et al.*, 2014; Wang *et al.*, 2014; Vogt-Eisele *et al.*, 2014; Yu *et al.*, 2017; Sacktor and Fenton, 2018). Although targeted candidate molecule approaches have shown that PKM*ζ* interacts with molecules that include kidney and brain protein (KIBRA), N-ethylmaleimide-sensitive factor (NSF), protein interacting with C-kinase 1 (PICK1), and the AMPA-type 2 glutamate receptor (GluA2). Although a picture has emerged from parallel studies that PKM*ζ* associates with KIBRA and activates NSF increasing the insertion of GluA2 subunit-containing *α*-amino-3-hydroxy-5-methyl-4-isoxazolepropionic acid (AMPA)-type glutamate ionotropic receptors into the postsynaptic density, coincident with reducing their PICK1-mediated endocytosis from the postsynaptic sites that are maintained by CaMKII-activity (Sacktor, 2011; Sacktor and Fenton, 2018), this hypothesis-driven approach is nonetheless for discovering and defining the network of interacting molecules that maintain memory. However, modern genomic techniques such as transcriptome profiling have made it possible to examine the molecular underpinnings of plasticity in animal behavior (reviewed in Harris and Hofmann, 2014). The present research combines hypothesis-driven techniques with the discovery-driven methods to identify novel neuromolecular substrates of memory and begin to reveal the network of molecular interactions that may underlie the maintenance of long-term memory.

In the present study we use an integrative approach across levels of biological organization to discover relationships between measures of behavior and gene expression. We generate a database by measuring mouse behavior on multiple dimensions of locomotion, learning and memory, and transcriptomes from tissue samples that were taken from the major hippocampal subfields after behaviorally evaluating long-term active place avoidance or null control memories. Specifically, our behavioral manipulations defined four behavioral groups: Two groups differed such that subjects could form one or two types of active place avoidance memory. The subjects in the other two yoked control groups could not form a memory for a place response, although the mice were exposed to the identical physical environment and stimuli. We used RNA-sequencing (RNA-seq) to examine the transcriptomes of three micro-dissected anatomically distinct subregions of the dorsal hippocampal circuit of the same subjects, the dentate gyrus, as well as the CA3 and CA1 subfields. We initially identified which genes were differentially expressed between the four behavioral groups and three subfields, focusing on the most variably expressed genes. We then examined whether gene expression for canonical memory-mediating molecules differed between the behavioral groups and the hippocampal subfields. Finally, we integrated across the levels of behavioral and transcriptomic biological data by harnessing the between-subject variation within each of the measurements. The co-variance structure between all pairs of measurements defined potential networks of high-order interactions across these levels of biological function. We then examined these interaction networks to discover novel associations between genes and the behavioral expression of memory and to evaluate the hypothesized roles of key memory-mediating molecules (CaMKII, the atypical PKCs, and GluA2).

## Methods

All animal care and use comply with the Public Health Service Policy on Humane Care and Use of Laboratory Animals and were approved by the New York University Animal Welfare Committee and the Marine Biological Laboratory IACUC. Male C57BL/6J mice were housed at the Marine Biological Laboratory on a 12:12 (light: dark) cycle with continuous access to food and water in home cages with up to five littermates.

### The Active Place Avoidance Task

To examine spatial learning and memory, we used a well-established active place avoidance paradigm. Littermates were randomly assigned to one of our treatment groups (standard-trained, n=8; standard-yoked, n=8; conflict-trained, n=9; conflict-yoked, n=9, *Figure 1*). All mice were exposed to nine 10-min trials in the active place avoidance arena. Mice were placed on an elevated circular 40-cm diameter arena made of parallel bars that rotated at 1 rpm. The arena wall was transparent and thus contained the mouse on the arena while allowing it to observe the environment. The location of the mouse in the arena was determined from an overhead digital video camera interfaced to a PC-controlled tracking system (Tracker, Bio-Signal Group Inc., Acton, MA). Trained mice in the active place avoidance task are conditioned to avoid the location of mild shocks (constant current 0.2 mA, 500 ms, 60 Hz) that can be localized by visual cues in the environment. Yoked-control mice are delivered the identical sequence of shocks that was received by a particular trained mouse (*Figure 1*, Suppl. Table 2), the difference being that for the yoked mice, the shocks cannot be avoided or localized to a portion of the environment. Mice are allowed to become familiar with walking on the rotating arena during a pretraining trial with no shock. Then each mouse received three training trails separated by a 2-h inter-trial interval. The mice were returned to their home cage overnight. The next day, each mouse received a “Retest trial” with the shock in the same location as before. For the next three training trials, the shock zone remains in the same place for standard-trained animals but is relocated 180° for the conflict-trained mice. The next day, all mice receive a memory “Retention trial” with the shock off to evaluate the strength of the conditioned avoidance.

**Figure 1.**
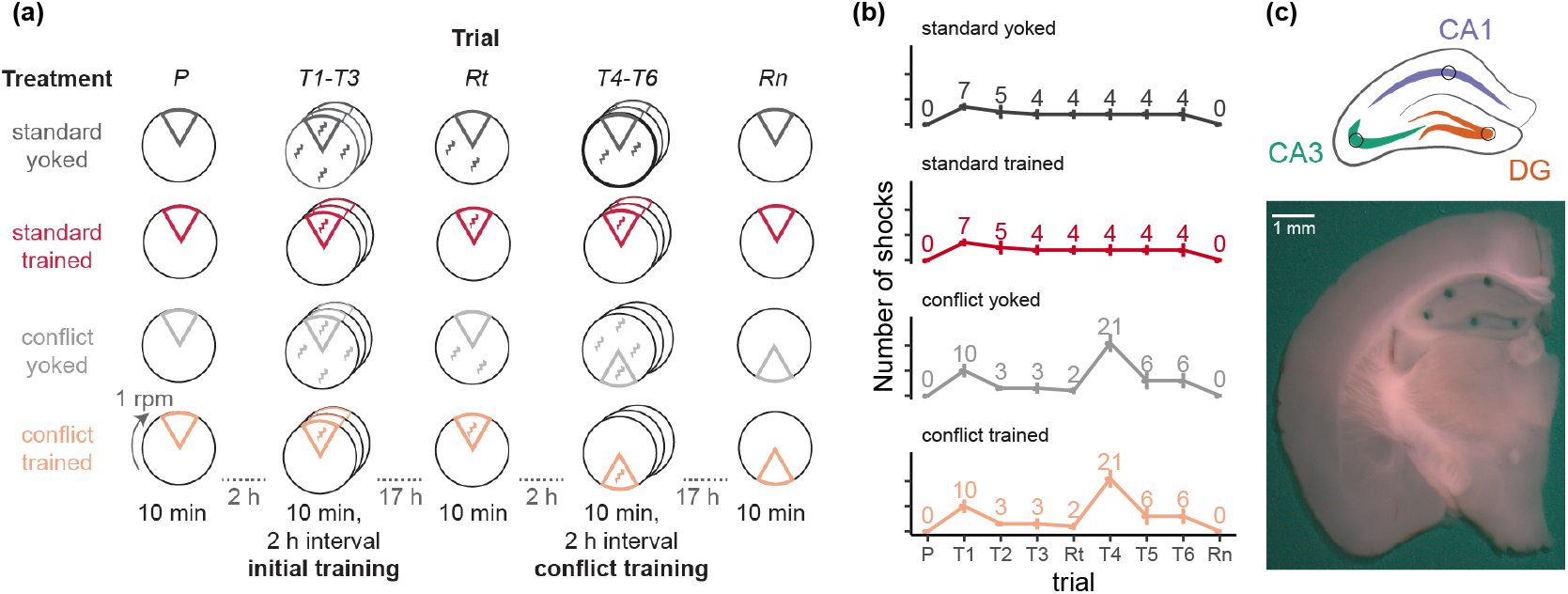
Experimental Design. **a)** Mice were assigned to one of four groups: standard-trained (red, n=8), standard-yoked (dark grey, n=8), conflict-trained (peach, n=9), or conflict-yoked (light grey, n=9). Mice were placed on the rotating arena (1 rpm) for trials that lasted 10 min and were separated by 2-h inter-trial intervals or overnight (~ 17 h). Behavior was recorded during the pretraining (P), initial training (T1-T3), retest (Rt), conflict training (T4-T6), and retention (Rn) trials. In the trial schematics, the 60° -sector represents the shock zone or for the yoked groups, the equivalent region that is used for evaluating place avoidance. Whereas the trained mice receive shocks only in the shock zone indicated by the sector, the yoked mice experience shock throughout the arena. **b)** Standard-trained and standard-yoked mice received an identical number of shocks (and time series), with the most shocks occurring on trial 1. Conflict-trained and conflict-yoked mice received an identical number of shocks (and time series), with the most shocks occurring on trial 4, the first trial of conflict training. **c)** A schematic representation and a photo show the size and location of tissue samples collected from the dorsal hippocampus for RNA-sequencing. Tissues from five subfields (CA1, CA2, CA3, CA4, DG) were collected, but only the DG (orange), CA3 (teal), and CA1 (purple) subfields were processed for RNA-seq.

### RNA-sequencing and bioinformatics

For RNA-sequencing, the DG, CA3, CA1 subfields were micro-dissected using a 0.25 mm punch (Electron Microscopy Systems) and a Zeiss dissecting scope (*Figure 1*). RNA was isolated using the Maxwell 16 LEV RNA Isolation Kit (Promega). RNA libraries were prepared by the Genomic Sequencing and Analysis Facility at the University of Texas at Austin and sequenced on the Illumina HiSeq platform. Reads were processed on the Stampede Cluster at the Texas Advanced Computing Facility. Quality of raw and filtered reads was checked using the program FASTQC (Wingett and Andrews, 2018) and visualized using MultiQC (Ewels *et al.*, 2016). We obtained 6.9 million ± 6.3 million reads per sample (*Figure S1*). Next, we used Kallisto to pseudo-align raw reads to a mouse references transcriptome (Gencode version 7 (Waterston *et al.*, 2002)), which yielded 2.6 million ± 2.1 million reads per sample (*Figure S1*). Mapping efficiency was about 42% (*Figure S1*). Transcript counts from Kallisto (Bray *et al.*, 2016) were imported into R (R Development Core Team, 2013) and aggregated to yield gene counts using the gene identifier from the Gencode transcriptome. DESeq2 was used to normalize and quantify gene expression with a false discovery corrected (FDR) p-value < 0.1 (Love, Huber and Anders, 2014; Morgan *et al.*, 2019). ShinyGo (Ge and Jung, 2018) was used to identify Gene Ontology terms associated with genes that are correlated with PC1. All genes associated with particular GO terms were identified using the Gene Ontology Browser (Krupke *et al.*, 2017; Bult *et al.*, 2019; Smith *et al.*, 2019). We compared the GO terms for candidate genes, differentially expressed genes, and a list of genes identified as important for long-term potentiation (Sanes and Lichtman, 1999; Harris *et al.*, 2019). We relied on the R packages ggplot2 (Wickham, 2009), cowplot (Wilke, 2016), and corrr (Ruiz, Jackson and Cimentada, 2019) for data visualization. Illustrations were created using Adobe Illustrator.

### Statistical analyses

Spatial behavior was evaluated by automatically computing (TrackAnalysis software (Bio-Signal Group Corp., Acton, MA) 26 measures that characterize a mouse’s use of space during the trial (**Supp. Table 1**). All statistical analyses were performed using R version 3.6.0 (2019-04-26) - “Planting of a Treeε (R Development Core Team, 2013), relying heavily on the software from the tidyverse library (Wickham, 2009; Wickham and Grolemund, 2017). Principal component analysis (PCA) was conducted to reduce the dimensionality of the data (Lê, Josse and Husson, 2008; Kassambara and Mundt, 2017). One- and two-way ANOVAs were used to identify group differences in behavioral measures across one or multiple trials, respectively (Stanley, 2018). For statistical analysis of gene expression, we used DESeq2 to normalize and quantify gene counts with a false discovery corrected (FDR) p-value < 0.1 (Love, Huber and Anders, 2014). DESeq2 models evaluated gene expression differences either between the four behavioral treatment groups (standard-trained, standard-yoked, conflict-trained, and conflict-yoked) or between the combined memory-trained and combined yoked-control groups.

### Data availability

Raw sequence data and differential gene expression data are available in NCBI’s Gene Expression Omnibus Database (accession:GSE99765 https://www.ncbi.nlm.nih.gov/geo/query/acc.cgi?acc=GSE99765). All data, code, and the results are publicly available on GitHub (https://github.com/raynamharris/IntegrativeProjectWT2015) with a stable version archived at Zenodo (Harris, 2017). Tables and supplementary tables are archived in Dryad (Harris *et al.*, 2020). An overview of the bioinformatic workflow used to create the figures and tables in the manuscript is described in **Suppl. Figure 2**.

## Results

### Characterizing cognitive behavior

We began by analyzing the place avoidance behavior amongst the memory-trained and corresponding yoked control groups, recalling that the corresponding training and control groups received the identical time series of shocks (*Figure 1*). Avoidance was evaluated in multiple ways, including as a reduction in the number of entrances to the shock zone, an increase in the time to first enter the shock zone, and a decreased likelihood of being in the shock zone (*Figure 2*). As expected, the memory-trained groups show avoidance behavior and the yoked groups do not show avoidance behavior, even though the corresponding trained and yoked control groups experience identical physical conditions. These observations were confirmed by two-way ANOVAs, with significant effects of training treatment, trial, and their interaction (see Table 1).

**Figure 2.**
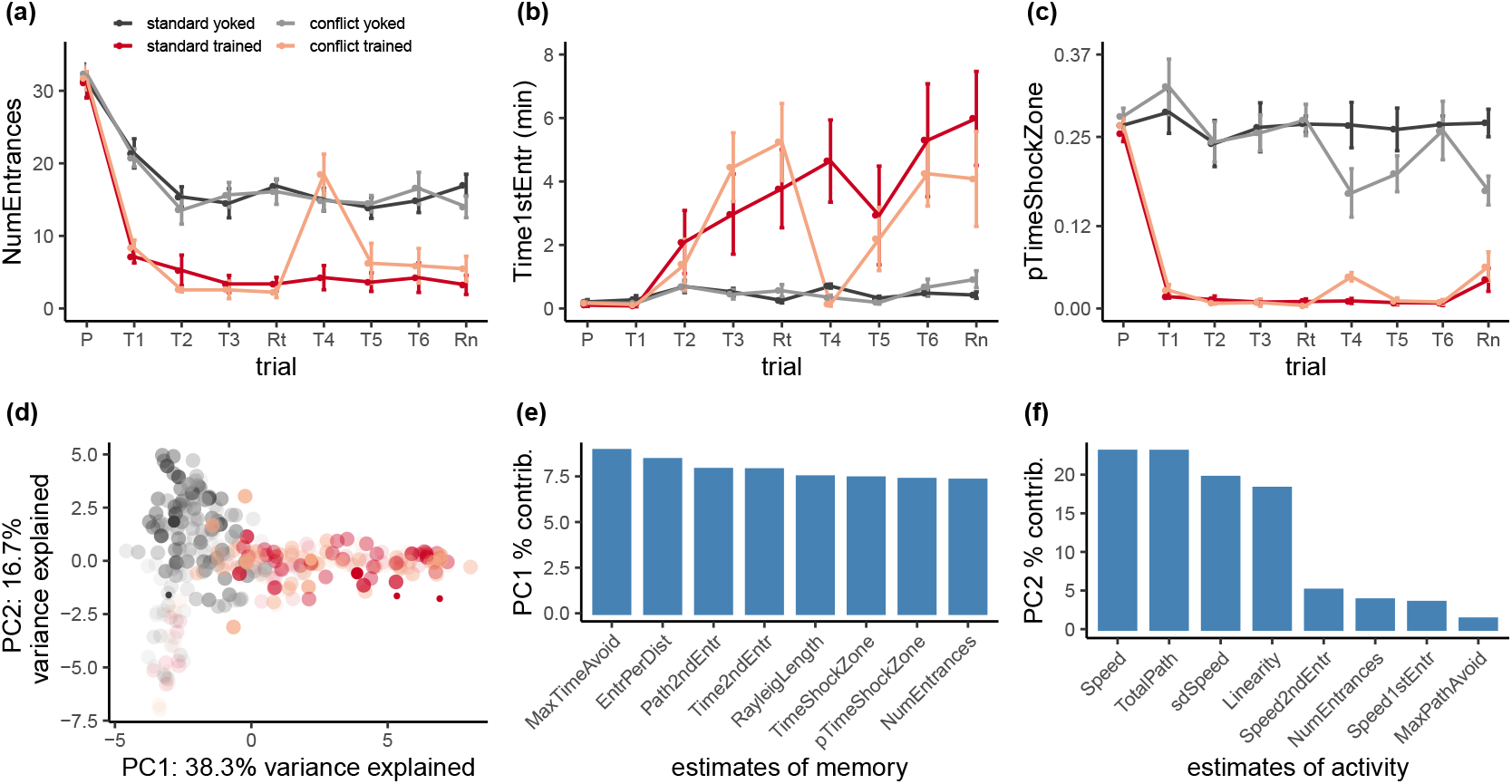
Characterizing and comparing behavior amongst the groups. Standard-trained and conflict-trained groups rapidly learned to effectively avoid the shock zone whereas the yoked control groups did not avoid the corresponding region measured by **a)** reduction in the number of entrances into the shock zone, **b)** and increased latency to enter the shock zone for the first time, **c)** and a reduction in the proportion of time spent in the shock zone. **d)** PCA performed on the 26 measures of behavior taken across all 9 trials, results in PC1 that distinguishes the trained and yoked groups, in particular during the retention test just prior to sacrifice when physical conditions are identical for all the mice because there is no shock. Large dots indicate group averages on the retention test while transparency is highest for pretraining and lowest for retention. **e,f)** Parameter loadings for PC1 and PC2 indicate that PC1 is comprised of estimates of place avoidance and PC2 estimates of locomotor behavior. Statistics are provided in Table 1. **Key:** NumEntrances = total number of entrances into the shock zone; Time1stEntr = time (min) passes before entering the shock zone the first time; pTimeShockZone = proportion of time spent in the shock zone relative to the total time spend in the four equivalent areas centered in each quadrant; MaxTimeAvoid = maximum time (s) avoiding the shock zone; TimeShockZone = total time (s) spent in the shock zone; SpeedArena.cm.s = average speed estimated in each 2 second epoch; TotalPath = total distance walked; Path2ndEntr = distance walked before entering the shock zone twice; EntPerDist = number of entrances per meter walked; MaxPathAvoid = maximum distance walked without entering the shock zone; Linearity = straightness of locomotion, estimated every 2 seconds as the ratio of the end-to-end Euclidean distance divided by the integral of the path segments; Speed1stEntr = speed when entering the shock zone the first time; Speed2ndEntr = speed when entering the shock zone the second time; sdSpeed = standard deviation of the speed estimates; RayleighLength = the Rayleigh vector length estimating the angular bias of the animal’s dwell probability.

**Table 1.** One-way ANOVAs were used to determine the effects of training treatment on behavior during a single trial, including the pretraining and recall trails. Two-way ANOVAs were used to determine the effects of treatment, training, and the interaction across three trials, including the initial training session and the conflict training session. Key: NumEntrances = total number of entrances into the shock zone; Time1stEntr = time (min) passes before entering the shock zone the first time; pTimeShockZone = proportion of time spent in the shock zone relative to the total time spend in the four equivalent areas centered in each quadrant.

| trials | ANOVA | Predictor | df | F | p | sig |
| --- | --- | --- | --- | --- | --- | --- |
| Pre-training (Pre) | NumEntrances ~treatment | treatment | 3, 30 | 0.09 | 0.967 |  |
| Pre-training (Pre) | pTimeShockZone ~treatment | treatment | 3, 30 | 0.78 | 0.512 |  |
| Pre-training (Pre) | Time1stEntr ~treatment | treatment | 3, 30 | 0.8 | 0.506 |  |
| Initial training (T1 - T3) | NumEntrances ~treatment * trial | treatment | 3, 90 | 26.42 | 0 | *** |
| Initial training (T1 - T3) | NumEntrances ~treatment * trial | trial | 2, 90 | 5.88 | 0.004 | ** |
| Initial training (T1 - T3) | NumEntrances ~treatment * trial | treatment x trial | 6, 90 | 0.72 | 0.631 |  |
| Initial training (T1 - T3) | pTimeShockZone ~treatment * trial | treatment | 3, 90 | 48.26 | 0 | *** |
| Initial training (T1 - T3) | pTimeShockZone ~treatment * trial | trial | 2, 90 | 0.88 | 0.419 |  |
| Initial training (T1 - T3) | pTimeShockZone ~treatment * trial | treatment x trial | 6, 90 | 0.59 | 0.736 |  |
| Initial training (T1 - T3) | Time1stEntr ~treatment * trial | treatment | 3, 90 | 0.02 | 0.997 |  |
| Initial training (T1 - T3) | Time1stEntr ~treatment * trial | trial | 2, 90 | 0.11 | 0.895 |  |
| Initial training (T1 - T3) | Time1stEntr ~treatment * trial | treatment x trial | 6, 90 | 3.24 | 0.006 | ** |
| Initial recall (Rt) | NumEntrances ~treatment | treatment | 3, 30 | 45.44 | 0 | *** |
| Initial recall (Rt) | pTimeShockZone ~treatment | treatment | 3, 30 | 129.49 | 0 | *** |
| Initial recall (Rt) | Time1stEntr ~treatment | treatment | 3, 30 | 7.96 | 0 | *** |
| Conflict training (T4 - T6) | NumEntrances ~treatment * trial | treatment | 3, 90 | 9.37 | 0 | *** |
| Conflict training (T4 - T6) | NumEntrances ~treatment * trial | trial | 2, 90 | 0.09 | 0.913 |  |
| Conflict training (T4 - T6) | NumEntrances ~treatment * trial | treatment x trial | 6, 90 | 3.43 | 0.004 | ** |
| Conflict training (T4 - T6) | pTimeShockZone ~treatment * trial | treatment | 3, 90 | 25.26 | 0 | *** |
| Conflict training (T4 - T6) | pTimeShockZone ~treatment * trial | trial | 2, 90 | 0.03 | 0.972 |  |
| Conflict training (T4 - T6) | pTimeShockZone ~treatment * trial | treatment x trial | 6, 90 | 1.54 | 0.174 |  |
| Conflict training (T4 - T6) | Time1stEntr ~treatment * trial | treatment | 3, 90 | 6.03 | 0.001 | ** |
| Conflict training (T4 - T6) | Time1stEntr ~treatment * trial | trial | 2, 90 | 0.05 | 0.954 |  |
| Conflict training (T4 - T6) | Time1stEntr ~treatment * trial | treatment x trial | 6, 90 | 1.74 | 0.12 |  |
| Conflict recall (Rn) | NumEntrances ~treatment | treatment | 3, 30 | 18.01 | 0 | *** |
| Conflict recall (Rn) | pTimeShockZone ~treatment | treatment | 3, 30 | 26.9 | 0 | *** |
| Conflict recall (Rn) | Time1stEntr ~treatment | treatment | 3, 30 | 5.97 | 0.003 | ** |
| All trials | PC1 ~treatment | treatment | 3, 302 | 91.83 | 0 | *** |
| All trials | PC2 ~treatment | treatment | 3, 302 | 10.76 | 0 | *** |
| Retention (Rn) | PC1 ~treatment | treatment | 3, 30 | 15.18 | 0 | *** |
| Retention | PC2 ~treatment | treatment | 3, 30 | 3.25 | 0.035 | * |

During the pretraining session, when the shock was off, there were no differences between the groups, as expected. Both the standard and conflict place avoidance trained groups showed rapid avoidance learning of the initial location of the shock location (T1-T3). Initial recall (or retest (Rt) of the conditioned place avoidance on the next day, also showed similar avoidance in the two trained groups, and as expected, avoidance was not observed in the yoked controls. Place avoidance in the two trained groups differed when the shock zone was subsequently relocated for the conflict group (T4-T6), specifically on the first trial with the new shock location (T4), as expected. This conflict trained group also rapidly learned to avoid the relocated shock zone on subsequent trials. In fact, on subsequent trials, their ability to avoid could not be distinguished from the group that received standard training to the initial shock location and demonstrating that compared to the standard-trained group, the conflict trained group learned to avoid more places. The ability to avoid the current location of shock was indistinguishable between the standard- and conflict- trained groups and behavior between the two yoked controls were also not distinct from each other. The day after the conditioning trials, on the memory retention test (Rn) with no shock, the conditioned avoidance persisted and was indistinguishable between the two trained groups but was not expressed in the yoked controls. These group similarities and differences reflect knowledge about the location of shock rather than the experience of shock *per se,* because the 24 hours before sacrifice were physically identical across the groups, constituting time in the home cage and 10 minutes on the arena with no shock.

Because spatial knowledge and memory are complex phenomena that we aim to relate to gene expression, we examined the possibility of describing the differences between the groups with a statistically valid metric that does not privilege one estimate of conditioned avoidance over another. After computing 26 measures from the trajectories during the behavioral sessions (**Suppl. Table 1**), we performed a principal component analysis on these data in an effort to reduce the dimensionality of describing different aspects of the measured behavior. PC1 explains 41 % of the variance and separates trained and yoked mice (*Figure 2*). Measurements related to shock avoidance and avoidance conditioning load most strongly on PC1 (*Figure 2*). In particular, as shown by the large dots in *Figure 2*, PC1 clearly distinguishes the trained and yoked groups in the absence of shock during the memory retention test just prior to sacrifice (p < 0.001, Table 1). PC2 explains 17% of the variance and is comprised of variables that describe locomotor activity (*Figure 2*) and also appear to distinguish the standard and conflict experiences of the mice (p < 0.05, Table 1). PC1 and PC2 were significantly negatively correlated (*Figure S3*). The sign of a PC is not informative as it was arbitrarily chosen, and therefore the sign of correlations to these PC estimates of memory and activity are also arbitrary. The meaning of any relationships must come from knowledge of the underlying biology, and we conclude that estimates of memory can be best summarized by PC1.

### Region-specific, cognitive behavior-driven differential gene expression

We next used RNA-seq to examine gene expression patterns in the DG, CA3, and CA1 subfields of the hippocampus in the different groups of mice on the hypotheses that differential gene expression in the hippocampus will be sensitive to spatial cognitive behavior and will differ across the DG, CA3, and CA1 subfields (*Figure 3*). Indeed, gene expression differed significantly amongst the groups and within the different subfields 24 hours after memory training and thirty minutes after the retention test during which the mice express active place avoidance memory without reinforcement. The lists of differentially expressed genes are provided in **Suppl. Table 4** and **Suppl. Table 5**.

**Figure 3.**
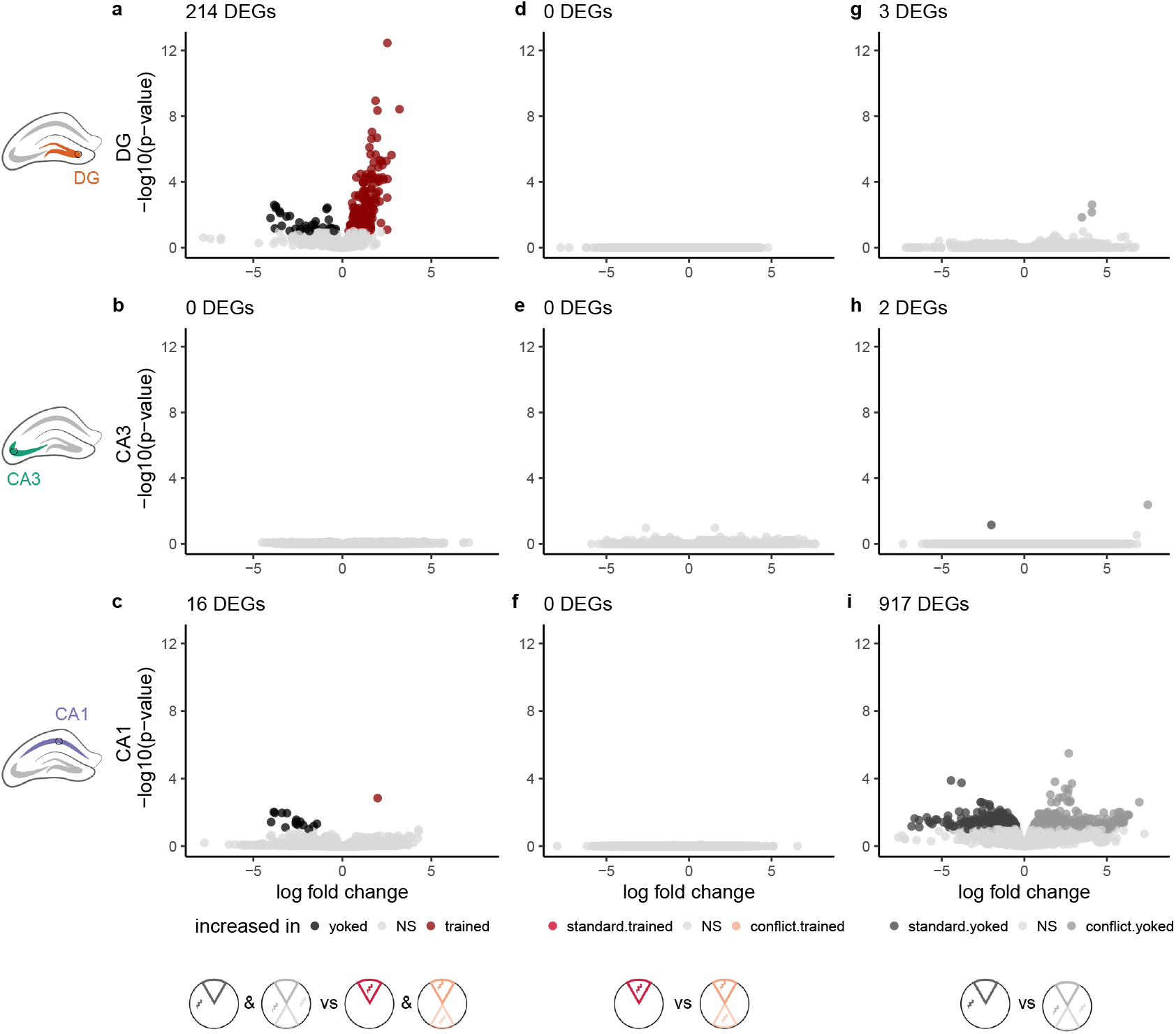
Subfield and treatment-specific gene expression differences. Volcano plots illustrating the significant and non-significant differences in gene expression when **a-c)** the two yoked groups are compared to the two trained groups, **d-e)** the two trained groups are compared, and **f-i)** the two yoked groups are compared. Rows show gene expression differences by region from DG **(a,d,g)** where information enters the hippocampus, to CA3 **(b,e,h)**, and CA1 **(c,f,i)**, the output of the hippocampus. Each dot represents a gene whose color corresponds to whether its expression increased, decreased, or was not significantly different (NS) between the two groups compared. The total number of differentially expressed genes (DEGs) are reported in the top left corner of each panel.

Because the behavior during the retention test did not differ between the standard-trained and conflict-trained groups, we combined the groups and compared all yoked to all trained mice (*Figure 3*). Gene expression differed (FDR p ≤ 0.1) between the trained and yoked groups the most in the DG, with substantially more increases of gene expression than decreases in the trained mice compared to the yoked controls (*Figure 3*). No differentially expressed genes were detected in CA3 and only a handful of genes were differentially expressed in CA1 following either standard or conflict training. Next we compared gene expression of standard-trained and conflict trained mice (*Figure 3*). We found no detectable changes in gene expression in DG, CA3, or CA1 even though mice in the conflict group received more shocks and learned two locations of shock (*Figure 3*).

Because the behavior during the retention test did not differ between the standard-trained and conflict-trained groups, we combined the groups and compared all yoked to all trained mice (*Figure 3*). Gene expression differed (FDR p ≤ 0.1) between the trained and yoked groups the most in the DG, with substantially more increases of gene expression than decreases in the trained mice compared to the yoked controls (*Figure 3*). No differentially expressed genes were detected in CA3 and only a handful of genes were differentially expressed in CA1 following either standard or conflict training (*Figure 3*). Next we compared gene expression of standard-trained and conflict trained mice (*Figure 3*). We found no detectable changes in gene expression in DG, CA3, or CA1 even though mice in the conflict group received more shocks and learned two locations of shock (*Figure 3*).

Finally, we compared the standard-yoked and conflict-yoked mice (*Figure 3*). Yoked mice receive exactly the same number and time series of shocks as their trained counterparts, but conflict-yoked mice received more shocks that standard-yoked mice. Thus, this comparison provides insight into how gene expression may change in the absence of a conditioned spatial memory. We found that differential gene expression in the standard-yoked and conflict-yoked groups was nearly absent in DG and CA3, but it is substantial in CA1 (*Figure 3*). In CA1, approximately equal numbers of genes were increased or decreased when the two yoked groups are compared (*Figure 3*). Together with the lack of differential gene expression between the two trained groups (*Figure 3*), this suggests that gene expression in CA1 is sensitive to unidentified, non-spatial features of experience, but not merely sensitive to the amount of shock.

### Molecules-associated with active place avoidance behavior

Since the DG was most sensitive to the place memory training, subsequent analyses focus on the 214 differentially expressed genes in DG between the combined trained and yoked control groups (**Suppl. Table 3**). This comparison is interesting because the physical environments did not differ between the trained and control groups (*Figure 2*), but what the animals could encode, learn, and remember did systematically differ between the two groups, as indicated by the presence and absence of robust place avoidance memory (*Figure 3*).

To test four hypotheses for how the trained and yoked control groups would differ, we identified four Gene Ontology terms in the Biological Process domain that map onto each hypothesis: 1) response to stimulus, 2) translation, 3) synapse organization, and 4) learning or memory (**Table 2, C1**). Next, we asked which genes in those GO domains are differentially expressed in the DG 24 hours after memory training and thirty minutes after the memory retention test (**Table 2, C2**). The majority of the genes that are differentially expressed in the DG are well known for their response to stimuli. Multiple immediate genes (e.g. *Arc, Fos, Fosl2, Egr1, Jun, Homer1*) and other transcription factors are differentially expressed in the DG. This is consistent with notion that immediate early gene expression is associated with the ongoing process of memory updating, not merely the formation of memory because further place avoidance learning was unlikely in the absence of reinforcement during the retention test response. *Neuro6d* was the only transcription factor that showed decreased expression, whereas *Npas4,* which is thought to mediate learning-related plasticity, especially of neural inhibition (Lin *et al.*, 2008; Spiegel *et al.*, 2014; Weng *et al.*, 2018) was more highly expressed in the DG. With respect to translation, we found that spatial training was only associated with increases in *Cpeb4* and *Eif5* expression. *Amigo2, Arc, Bdnf, Flrt3, Fzd5, Homer1, Lrrtm2, Npas4, Pcdh8,* and *Slitrk5* are all important for synapse organization and were increased in the trained DG. Fifteen genes from the “learning or memory” GO domain were found to be differentially expressed; all of these genes appear in one or more of the other three GO domains. To estimate whether the results would differ if we chose arbitrary GO terms we randomly selected the following terms: epithelial cell proliferation (GO:0050673), reproductive process (GO:0022414), rhythmic process (GO:0048511), and multi-organism process (GO:0051704) and find that 18 DEGs were among the 4273 (0.41 %) total genes in the random GO terms compared to the 90 DEGs that were among the 7156 (1.48%) genes in our hypothesized GO terms (proportions test z = 10.92; p = 10^23^), indicating that more differentially expressed genes were part of the hypothesized GO categories than would be expected by chance.

**Table 2.** Analysis of GO terms hypothesized to play a role in learning and memory. Column 1 contains four GO categories representing four hypotheses about which genes are important for memory formation and recall: 1) response to stimulus, 2) translation, 3) synapse organization, and 4) learning or memory. Column 2 contains genes that were differentially expressed in the dentate gyrus (DG) of trained and yoked control mice a day after memory training and thirty minutes after unreinforced expression of active place avoidance memory. Of those DEG, 79 are associated with hypothesis 1 that genes would be related to response to stimulus, which includes many immediate early genes that trigger responses in signaling cascades with membrane-bound receptors. Only two genes were associated with the hypothesis that translation machinery that executes the response are needed. Only ten genes are associated with “synaptic organization” that includes synaptic receptors and channels that express the response. Fifteen genes associated with learning or memory at the behavioral level were differentially expressed. *Adrb1,Arc, Bdnf, Btg2, Egr1* (aka ZIF-268), *Hmgcr, Jun, Npas4, Nptx2, Pak6, Plk2, Ptgs2, Sgk1, Srf,* and *Syt4* All differentially expressed genes in the learning and memory list also overlap with the genes in the response to stimulus list. Column 3 contains the candidate genes described in the introduction or discussion and includes *Camk2a, Fmr1, Gria2, Igf2, Mtor, Pick1, Prkcz* (PKM *f*), and *Wwc1* (KIBRA). Column 4 contains genes that were reviewed by Sanes and Lichtman, 1999 to play a role in long-term potentiation, a cellular model of memory. Only four genes (shown in bold: *Adrb1, Bdnf, Egr1,* and *Homer1)* from the Sanes and Lichtman list were found to be differentially expressed in the DG.

| GO terms | DG DEGs | Candidates | LTP genes |
| --- | --- | --- | --- |
| Response to stimulus (GO:0050896) | *Abhd2, Adrb1, Ahr, Ankrd27, Apaf1, Arc, Arid5b, Arl13b, Arpp21, Atf3, B3gnt2, Bach1, BC048403, Bmt2, Btg2, C2cd4b, Ccnk, Cited2, Fbxw7, Fermt2, Flrt3, Fos, Fosb, Fosl2, Foxg1, Foxo1, Fzd4, Fzd5, Gadd45g, Gnaz, Gpi1, Gpr19, Hmgcr, Homer1, Hspa1a, Hsph1, Il16, Ing2, Irs1, Irs2, Jun, Junb, Jund, Kdm6b, Kitl, Klf2, Klf6, Klkb1, Lbh, Lemd3, Lmna, Mc1r, Mest, Myc, Nedd9, Nfil3, Npas4, Nptx2, Nr4a1, Nr4a2, Nr4a3, Ppp1r15a, Slc16a1, Slc25a25, Slitrk5, Smad7, Sox9, Srf, Srgap1, Stac2, Syt4, Thbs1, Tiparp, Tnip2, Tra2b, Trib1, Tsc22d2, Zbtb33, Zfand5* | *Camk2a, Fmr1, Gria2, Igf2, Mtor, Wwc1* | *Adcy1, Adra2a, Adra2b, Adra2c, Adrb1, Adrb2, Adrb3, Cacna1a, Cacna1b, Cacna1c, Cacna1d, Cacna1e, Cacna1f, Cacna1s, Calb1, Calm1, Calm2, Calm3, Camk1, Camk4, Capn1, Capn10, Capn2, Capn3, Ccr7, Cd47, Cdh1, Cdh2, Chrm1, Chrm2, Chrm3, Chrm4, Chrm5, Chrna1, Chrna3, Chrna7, Chrnb1, Chrnb2, Chrnb3, Cnga2, Fgf2, Fyn, Gabbr1, Gabra1, Gabra2, Gabra3, Gabra5, Gabra6, Gabrb1, Gabrb2, Gabrb3, Gabrr1, Gap43, Gfap, Gria1, Gria2, Grin1, Grin2a, Grin2d, Grm1, Grm4, Grm5, Grm7, Gucy1a2, Gucy1b2, Gucy2c, Gucy2d, Gucy2e, Gucy2g, Homer1, Homer2, Homer3, Htr1a, Htr1b, Htr1f, Htr2a, Htr2b, Htr2c, Htr3a, Htr3b, Htr4, Htr5a, Htr5b, Htr6, Htr7, Il1b, Inhba, Itga1, Itga10, Itga11, Itga2, Itga2b, Itga3, Itga4, Itga5, Itga6, Itga7, Itga8, Itga9, Itgad, Itgae, Itgal, Itgam, Itgav, Itgax, Itgb1, Itgb1bp1, Itgb2, Itgb2l, Itgb3, Itgb4, Itgb5, Itgb6, Itgb7, Itgb8, Itgbl1, Itpkb, L1cam, Mapk1, Mapk11, Mapk12, Mapk14, Mapk3, Mapk4, Mapk6, Mapk7, Mapk8, Mapk9, Mas1, Ncam1, Ngf, Nos1, Nos3, Nrg1, Nrg2, Nrg3, Nrgn, Pnoc, Sptbn1, Src, Stx1b, Syp, Th, Thy1, Tnc, Ube3a, Vamp2, Vamp3, Vamp4, Vamp8* |
| Translation (GO:0006412) | *Cpeb4, Eif5* | *Fmr1, Mtor* | NA |
| Synapse organization (GO:0050808) | *Amigo2, Arc, Bdnf, Flrt3, Fzd5, Homer1, Lrrtm2, Npas4, Pcdh8, Slitrk5* | *Fmr1, Pick1* | *Ache, Bdnf, Cacna1a, Cacna1s, Camk1, Cdh1, Cdh2, Chrna1, Chrna7, Chrnb1, Chrnb2, Dlg4, Drd1, Efna5, EphA5, Erbb4, Fyn, Gabra1, Gabrb2, Gabrb3, Gap43, Grin1, Grin2a, Grm5, Homer1, Htr1a, Itga3, Itgam, Itpka, L1cam, Mapk14, Nrg1, Nrg2, Ntrk2, Ptn, Rab3a, Syn1, Tnc, Ube3a* |
| Learning or memory (GO:0007611) | *Adrb1, Arc, Bdnf, Btg2, Egr1, Hmgcr, Jun, Npas4, Nptx2, Pak6, Plk2, Ptgs2, Sgk1, Srf, Syt4* | *Igf2, Mtor, Prkcz* | *Adcy1, Adrb1, Adrb2, Bdnf, Cacna1c, Cacna1e, Calb1, Camk4, Chrna7, Chrnb2, Cnr1, Creb1, Drd1, Egr1, Gabra5, Gria1, Grin1, Grin2a, Grm4, Grm5, Grm7, Htr2a, Htr6, Htr7, Il1b, Itga3, Itga5, Itga8, Itgb1, Ncam1, Ngf, Ntrk2, Oprl1, Pla2g6, Plcb1, Prkar1b, Prkcz, Ptn, S100b, Snap25, Th* |

*Arc* is widely used as a marker of neural activity, so we were not surprised to find that *Arc* expression was positively correlated with PC1 (*Figure 4*), which suggests it is related to the persistence of conditioned avoidance memory. This training-induced relationship was not observed in the CA3 (r = −0.30, p = 0.327) or CA1 tissue (r = 0.20, p = 0.48) (*Figure 4*), nor were Arc and other immediate early genes differentially expressed in these samples suggesting the relationship has to do more with the memory-related information processing that each subregion is thought to be specialized for, rather than locomotor activity that is likely to be globally related to the hippocampus but a confound of avoidance memory expression. We also wanted to discover potential relationships between the persistence of conditioned avoidance behavior and gene expression in the DG in an unbiased manner, without selecting genes based on presumed knowledge and hypotheses. To visualize the relationship between these genes and behavior, we selected the 20 genes that were most positively correlated with PC1 (*Figure 4*). Then, we created a correlation network between all pairwise comparisons greater than the absolute value of 0.6 (*Figure 4*), which illustrates that all the genes that are positively correlated with PC1 are also positively correlated with one another. All immediate early genes are strongly positively correlated with each other and with PC1. Even with sample sizes of 16, many of the genes that are correlated are also differentially expressed and *vice versa.* This network of pairwise covariation amongst the differentially expressed genes suggest, as in the case of the immediate early genes, that there functionally related groups of genes can be identified by analysis of such covariance networks. To further explore potential responses to increased immediate early gene activity, we identified the top 5 GO terms associated with 85 genes that are both correlated with behavioral PC1 and differentially expressed in the DG (**Table 3**.) This data-driven analysis suggests that the molecular function of these genes is to regulate transcription via RNA pol-II activity in the dendritic regions of neurons to modulate the learning and memory.

**Figure 4.**
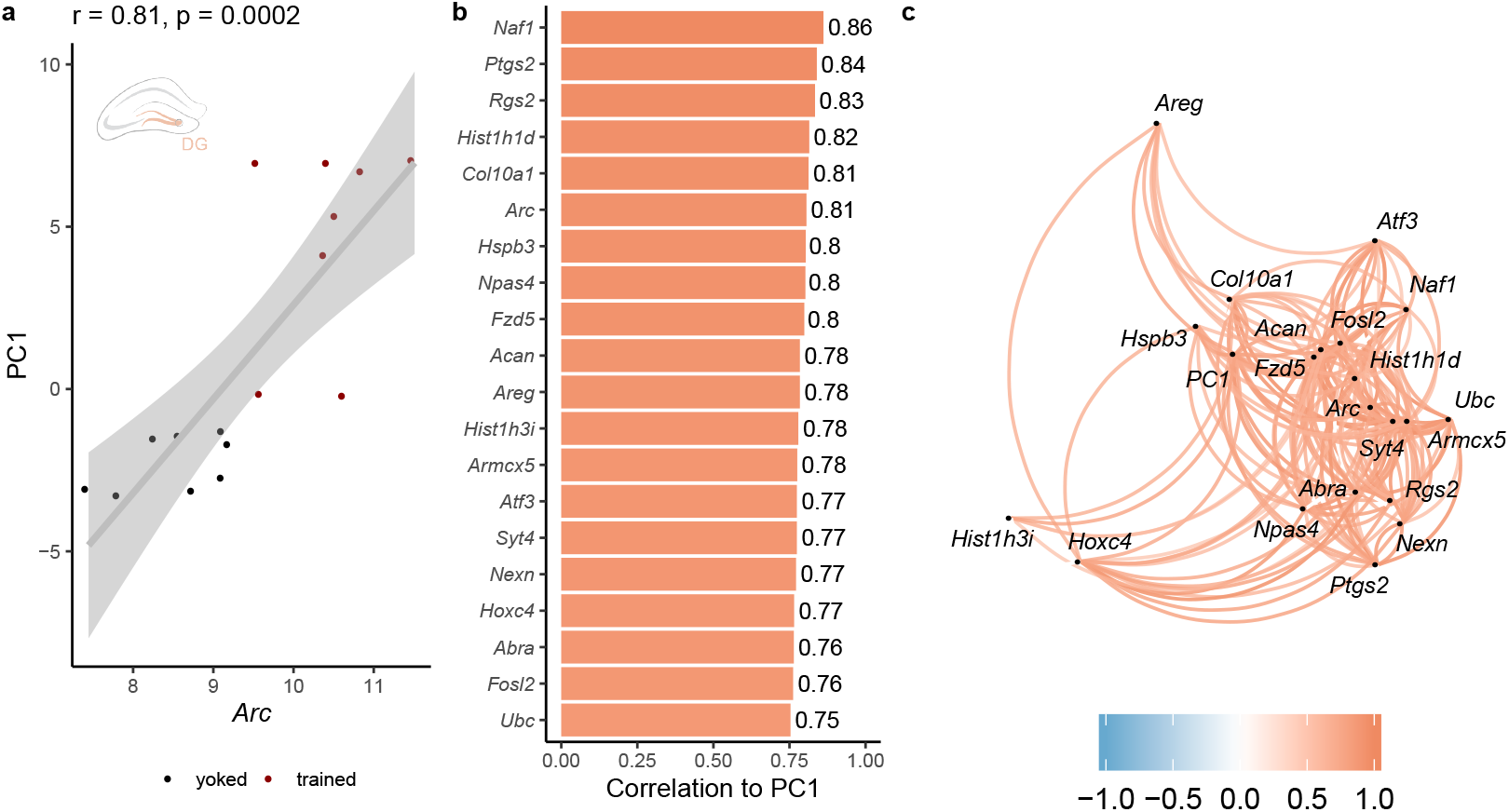
Correlations between gene expression in the DG and estimates of long-term avoidance memory. **a)** The immediate early gene Arc is differentially expressed in the DG and is positively correlated with behavior PC1 (an estimate of avoidance memory): red = trained, black = yoked. **c)** Arc is also among the top 20 genes that are positively correlated with PC1. For the full list of genes that are positively correlated with PC1, see Suppl. table 4. **d)** The genes that are positively associated with PC1 are highly correlated with each other. All correlations greater than 0.6 or less than −0.6 are shown in red or blue, respectively.

**Table 3.** Discovery-driven analysis of GO terms that are positively correlated with PC1. Column 1 lists the three GO domains: Biological Process (BP), Cellular Compartment (CC), and Molecular Function (MF). Columns 2 lists the top 5 most significant functional categories that are represented in the list of 58 genes that are positively correlated with PC1, a measure of avoidance behavior

| Domain | Functional Category | Gene |
| --- | --- | --- |
| BP | Memory | *Arc Bdnf Egr1 Kcnk10 Npas4 Plk2 Ptgs2 Sgk1 Syt4* |
| BP | Learning or memory | *Arc Bdnf Btg2 Egr1 Kcnk10 Npas4 Plk2 Ptgs2 Sgk1 Syt4* |
| BP | Tissue development | *Acan Arc Areg Atf3 Bdnf Btg2 Col10a1 Egr1 Errfi1 Fosl2 Frmd6 Fzd5 Homer1 Hoxc4 Nr4a3 Pcdh8 Ptgs2 Rgs2 Slc25a25 Smad7 Tiparp* |
| BP | Cognition | *Arc Bdnf Btg2 Egr1 Kcnk10 Npas4 Plk2 Ptgs2 Sgk1 Syt4* |
| BP | Behavior | *Arc Bdnf Btg2 Egr1 Homer1 Kcnk10 Npas4 Nr4a3 Plk2 Ptgs2 Sgk1 Slc16a1 Syt4* |
| CC | Neuron projection | *Acan Arc Bdnf Cpeb4 Fzd5 Homer1 Nexn Pcdh8 Plk2 Ptgs2 Rgs2 Sgk1 Syt4* |
| CC | Cell junction | *Arc Cpeb4 Frmd6 Fzd5 Homer1 Nexn Pcdh8 Slc16a1 Smad7 Syt4* |
| CC | Dendrite | *Arc Bdnf Cpeb4 Fzd5 Homer1 Pcdh8 Plk2 Syt4* |
| CC | Dendritic tree | *Arc Bdnf Cpeb4 Fzd5 Homer1 Pcdh8 Plk2 Syt4* |
| CC | Somatodendritic compartment | *Acan Arc Bdnf Cpeb4 Fzd5 Homer1 Pcdh8 Plk2 Syt4* |
| MF | Regulatory region nucleic acid binding | *Atf3 Egr1 Egr4 Fosl2 Hoxc4 Nfil3 Npas4 Nr4a3 Per1 Smad7 Tiparp* |
| MF | Transcription regulatory region DNA binding | *Atf3 Egr1 Egr4 Fosl2 Hoxc4 Nfil3 Npas4 Nr4a3 Per1 Smad7 Tiparp* |
| MF | RNA polymerase II regulatory region sequence-specific DNA binding | *Atf3 Egr1 Egr4 Fosl2 Hoxc4 Nfil3 Npas4 Nr4a3 Per1* |
| MF | DNA-binding transcription factor activity, RNA polymerase II-specific | *Atf3 Btg2 Egr1 Egr4 Fosl2 Hoxc4 Nfil3 Npas4 Nr4a3* |
| MF | RNA polymerase II regulatory region DNA binding | *Atf3 Egr1 Egr4 Fosl2 Hoxc4 Nfil3 Npas4 Nr4a3 Per1* |

Because PKM*ζ* is crucial for the persistence of active place avoidance memory in hippocampus (Pastalkova *et al.*, 2006; Sacktor, 2011)), we next examined several candidate genes in the PKM*ζ* signaling cascade. We also included *Igf2* because it promotes acquisition of memory, likely by an independent process (Chen *et al.*, 2011; Alberini and Chen, 2012). PKM*ζ* associates with KIBRA and activates NSF increasing the insertion of GluA2 subunit-containing *α* -amino-3-hydroxy-5-methyl-4-isoxazolepropionic acid (AMPA)-type glutamate ionotropic receptors into the postsynaptic density, coincident with reducing their PICK1-mediated endocytosis from the postsynaptic sites that are maintained by CaMKII-activity. Consequently, one might therefore predict significant negative and positive correlations in the expression of these genes on the hypothesis that their levels of transcription is a straightforward determinant of their function because PICK1 function is anti-related to the memory-promoting cell biological functions of PKM*ζ*, KIBRA, NSF, CaMKII and GluA2. Furthermore, LTP-induction is associated with transient upregulation of PKC *ι/λ*, the other atypical PKC, whereas genetic deletion of the PKM*ζ* gene Prkcz, results in LTP- and memory-triggered prolongation of compensatory upregulation of PKC*ι/λ* and the conventional isoform PKC*β*2 (Tsokas *et al.*, 2016). Consequently, on the same hypothesis, one might expect these molecules to also be positively related to each other and measures of memory. We used this hypothesis-driven candidate gene approach to select these genes and correlated their expression with the composite avoidance estimate expressed by PC1 (see above) for the three hippocampal subfields (*Figure 5*). As can be seen, *Igf2* expression was the most strongly positively correlated with the PC1 estimate of memory in each of the hippocampus subfields, but only the correlation between CA1 *Igf2* and PC1 reached significance. Of the memory maintaining candidate molecules, only the level of *Gria2* in CA3 was significantly correlated with PC1, but this was not predicted as it is a negative correlation. Because GluA2 mediates late-LTP, one would have expected the opposite relationship of increased *Gria2* expression to correlate with increased expression of memory. The correlations amongst the candidate genes tended to be weak, although there were some significantly strong relationships, less so in dentate gyrus and CA1 and more so in CA1 (*Figure 5*). The notion that the atypical PKCs may play complementary but competing roles in maintenance of LTP and memory, perhaps through binding to KIBRA, predicts anti-correlations between *Prkcz* and *Prkci* and whichever isoform is more expressed, to be correlated with KIBRA. This pattern was however not observed, suggesting a simple model of transcriptional regulation does not map their functional relationships.

**Figure 5.**
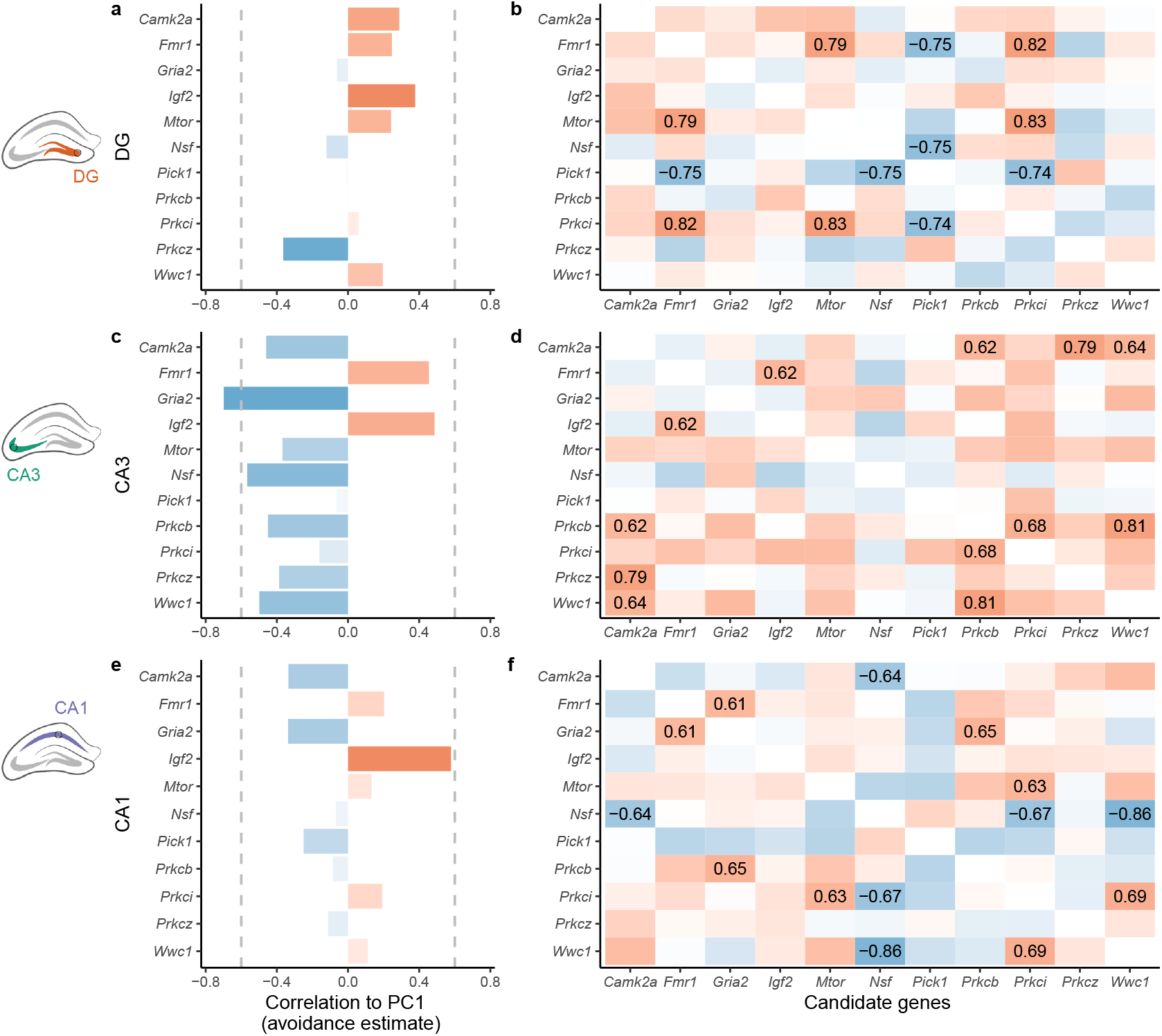
Region-specific candidate gene analysis of PKM*ζ* -related signaling genes. **a)** In the DG, none of the candidate memory-related genes are significantly correlated with behavioral PC1; however, **b)** *Pick1* is negatively correlated with *Fmr1, Nsf* and *Prkci.* The translation regulators *Fmr1* are *mTor* are positively correlated with each other. **c)** In CA3, *Gria2* is negatively correlated with PC1 and positively correlated with PC2. **d)** *Camk2a, Prkci, Prkcz,* and *Wwc1* all show positive correlations with one another. *Igf2* and *Fmr1* are also positively correlated. **e)** In CA1, *Igf2* is positively correlated with PC1. **f)** Nsf is negatively correlated with *Cam2ka, Prkci,* and Wwc7. *Gria2* is positively correlated with *Fmr1* and *Prkcb. Prkci* is positively correlated with Wwc7. Correlations greater than 0.6 or less than −0.6 are highlighted in the figure.

Finally, because glia and neurons contribute an approximately equal number of cells in our samples (Herculano-Houzel *et al.*, 2007), we considered several candidate genes associated with astrocytes, on the evidence that astrocytes contribute to memory. *Figure 6* illustrates that unlike the memory-associated candidate genes, not only were astrocyte-associated genes differentially expressed in the trained compared to the control samples, but in addition, many of their expression values in CA3 tended to co-vary with the PC1 estimate of memory. This was not the case in the dentate or CA1 samples. The expression of these genes was characteristically positively correlated, especially in CA3, but also in the dentate gyrus and to a lesser extent in CA1, which may in a sense, may at least partially reflect that we chose them because of their known association with astrocytic biology.

**Figure 6.**
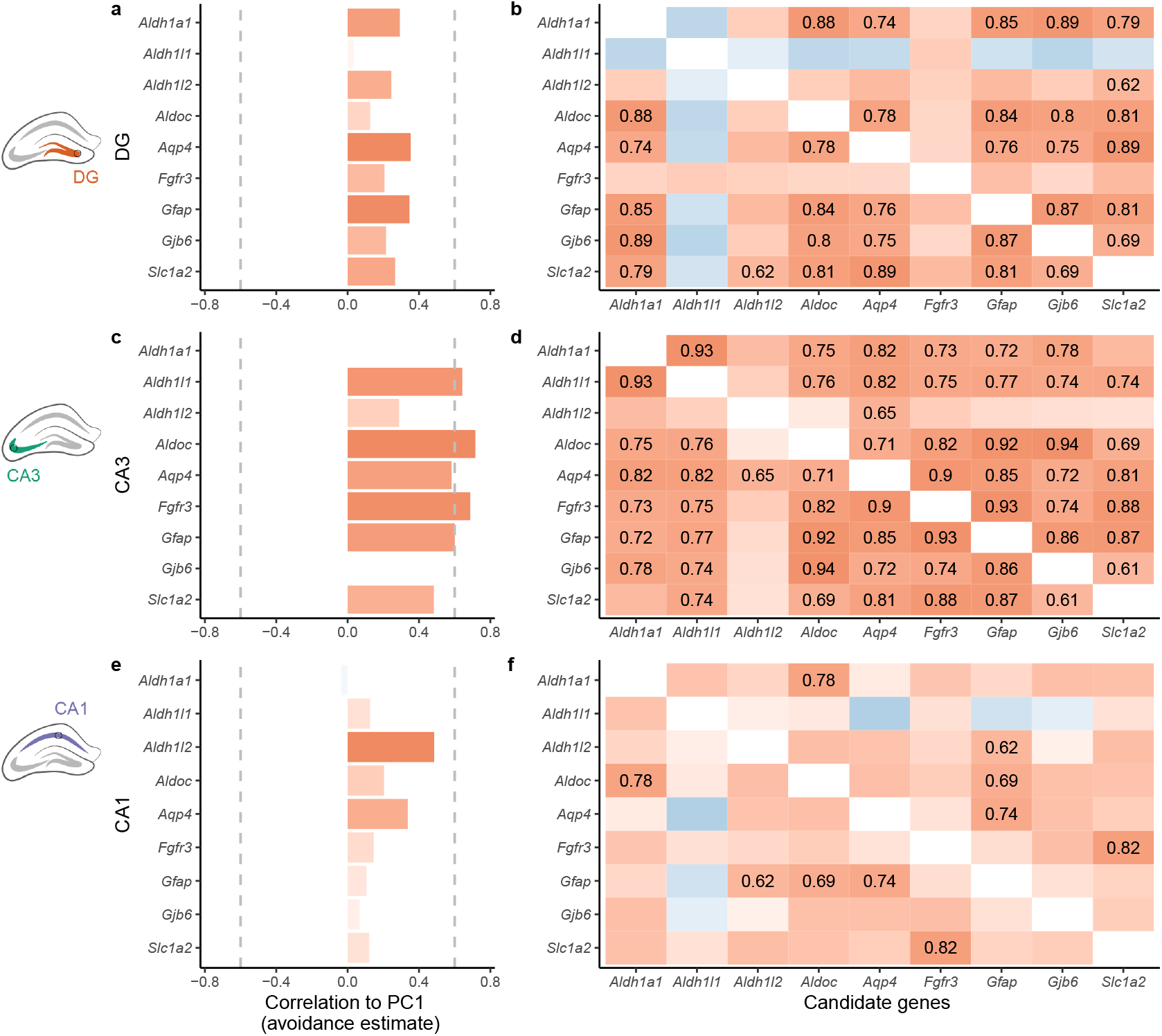
Region-specific candidate gene analysis of astrocyte-related genes. None of the astrocyte-related genes were significantly correlated in DG **(a)** or CA1 **(e)**, but *Aldh1l1, Aldoc, Fgfr,* and *Gfap* expression in the CA3 all show a correlations greater than 0.6 to behavioral PC1 **(c)**. These candidate genes are highly, positively correlated with one another, especially in the CA3 **(d)** and to a lesser extent in the DG **(b)** and CA1**(f)**.

## Discussion

In the present study we evaluated conditioned active place avoidance, and in particular long-term memory for the conditioned response by quantifying both the conditioned behavior and the associated transcriptional changes in three subfields of the hippocampus 24 hours later the end of the training and 30-min after expression of the avoidance memory. We first used an unbiased data-driven approach to identify genes that are sensitive to the maintenance of spatial memories across at least 24 h, and we used Gene Ontology analyses to identify the potential mechanisms for that these differentially expressed genes might have influenced brain structure and function. Then, we conducted a hypothesis-driven analysis of the data to characterize whether the memory training is associated with altered expression of dozens of genes that have been previously shown to be important for learning, memory, and the cellular model LTP. For this hypothesis-driven approach, we examined both differential gene expression and covariance with estimates of avoidance behavior and with other candidate genes. Our RNA-seq methods could not detect ribosomal RNA, nor did we perform protein sequencing, limiting the scope of this study to findings that may related to memory-related transcriptional regulation.

As expected, and by design, both memory-trained groups (standard and conflict) show avoidance behavior but their yoked counter parts do not. Because the comparison groups experience the identical physical conditions this design allows us to identify molecular responses to avoidance training and memory recall. Gene expression differs between the trained and yoked groups the most in the DG, with substantially more increases of gene expression than decreases in the trained mice compared to the yoked controls. Because the two trained groups did not differ, gene expression in DG is preferentially sensitive to active place learning, and spatial cognition in general, rather than to the specific number of places at which the mice were conditioned. Very few genes were differentially expressed in Ammon’s horn, which is expected if memory maintenance involves only a select number of molecular components, but is not expected in light of theories of hippocampus memory and information processing (Marr, 1971; Treves and Rolls, 1992; Lisman *et al.*, 2017) and findings that maintaining the active place avoidance memory persistently enhances CA3-CA1 synaptic function for at least 30-days (Pavlowsky *et al.*, 2017).

Nor did we find that memory-related differential expression of genes for the molecular translation machinery in the MEK/ERK and MTOR pathways. The majority of the genes that are differentially expressed in DG or are positively correlated with avoidance behavior belonged to transcription factors that regulate RNA polymerase II. Among these genes are the well-known immediate early genes Arc, Fos, and Jun, and Npas4. In addition to being differentially expressed, expression of these genes is highly correlated with one another. In previous experiments, Arc was activated during contextual threat avoidance conditioning and optogenetically silencing the Arc-activated DG neurons was sufficient to impair contextual fear memory, whereas optogenetically stimulating neurons that activated c-Fos was sufficient for expressing the memory (Liu *et al.*, 2012). Indeed, as we also observed in DG, prior work has shown that *Arc* and *Fos* expression are correlated in CA1 neurons regulating learning (Mahringer *et al.*, 2019). We cannot assert that the gene expression findings associated with active place avoidance memory are due to altered gene expression specifically in neurons because we sampled bulk tissue, containing approximately equal numbers of neurons and other cells that are largely the glial population (Herculano-Houzel *et al.*, 2007). Because we examined gene expression a day after training when training-associated immediate early gene activation is not expected, it is possible that the robust immediate early gene activation that we observed is related to the mice learning about the absence of shock during the retention test. However, that possibility predicts an inverse relationship between PC1 and immediate early gene expression because mice that expressed the weakest place avoidance by entering the shock zone the most during the retention test (low PC1 values) would have the greatest opportunity to learn that shock was absent and the 5 mice that showed perfect avoidance by never entering during the retention trial could not have learned that the shock was off. We reject this possibility because the correlations between immediate early gene expression in DG and PC1 on the retention test are all positive as is the correlation between Igf2 expression in DG, CA3, and CA1 with the PC1 estimate of memory (**Suppl. Fig 3d-f**). IGF-II activity promotes memory formation, enhancing the consolidation of long-term memory (Chen *et al.*, 2011), which is a process that can last days and relies on C/EPB signaling, that is likely related to the Cepb differential gene expression observed we observed in DG (Alberini *et al.*, 1994). Instead, these findings suggest that immediate early genes can respond not only to the novel experience of stimuli that are encoded during the conditioning of a response, but that their activation in DG also corresponds to the expression of long-term memory.

An analysis of the gene ontology revealed that many of the genes that are differentially expressed in the DG exert their function in dendrites and are important for neuronal projections and cell-cell junctions, including synapses, which is expected from the relationship between LTP and memory that forms the basis of the widely-accepted synaptic plasticity and memory hypothesis (Takeuchi, Duszkiewicz and Morris, 2014). Although, we cannot distinguish the contributions of neurons and glia to the memory-related differential gene expression we observed, these findings suggest that the expression of the conditioned place avoidance altered neural activity in a way that increases transcriptional activity, perhaps for the purpose of promoting dendritic growth. Indeed, we note that expression of a number of astrocyte-related genes in DG, CA3, and CA1 were positively correlated with the PC1 estimate of memory expression, including *Aldh1a1, Aldh1l2, Gfap, Fgfr3, and Aqp4* (**Fig. 6**). Even though none of these genes were differentially expressed after avoidance training, the correlation with expression of the conditioned avoidance behavior adds to the growing evidence that astrocytes also play an active role in forming and/or stabilizing memory, perhaps by calcium-dependent signaling at NMDA receptors (Parpura *et al.*, 1994; Suzuki and Shimodaira, 2006; Henneberger *et al.*, 2010; Suzuki *et al.*, 2011; Rusakov, 2015; Gao *et al.*, 2016).

Synaptic reorganization requires that neurons polarize their dendrites to create new neural connections and direct channels to specific regions within the synapse. Distruption of the processes that that regulate synaptic organization can impair memory or cause disease. Dysregulation of *Eif5* is associated with many neurodegenerative diseases (Kapur, Monaghan and Ackerman, 2017). *Cpeb4* encodes cytoplasmic polyadenylation element binding protein 4, which regulates the translation of genes by modulating their poly(A)-tail length and is a known regulator of genes associated with autism spectrum disorder (Parras *et al.*, 2018). Many of the genes that are mutated in autism spectrum disorder are crucial components of the activity-dependent signaling networks that regulate synapse development and plasticity (Ebert and Greenberg, 2013). Deletion of *Fmr1* can lead to dysregulated mRNA translation, altered synaptic function, loss of protein synthesis-dependent synaptic plasticity (Bassell and Warren, 2008; Thomson *et al.*, 2017; Talbot *et al.*, 2018). Some memory molecules have been characterized because they were found to be relevant in cancer, thus, it’s possible that studies of memory can lead to discoveries that are important for cancer or development.

Very few of the differentially expressed genes in our study overlapped with the LTP-related molecules that are identified from review of the LTP literature. Some classes of genes that are notably absent from the list of differentially expressed genes but present in the LTP-related list are genes for receptors (calcium, glutamate, GABA, and serotonin), calcium or calmodulin-binding proteins, and adhesion molecules. This is perhaps not surprising because the review focused on literature examining LTP induction, the first minutes of long-term potentiation, whereas our experiments were designed to investigate 24-h memory expression, which would be expected to correspond to LTP maintenance. Thus this work is consistent with the evidence that like LTP-induction, many molecules known to be crucial for learning, are neither crucial nor expressed in LTP maintenance and as we observe, memory persistence. This points to different molecular mechanisms for forming a memory and maintaining and/or later expressing it days later.

What should one then make of the atypical PKCs and CaMKII, candidate molecules that are demonstrated to have crucial roles in the maintenance of LTP and synaptic structure, as well as active place avoidance memory (Tsokas et al., 2016; Rossetti et al., 2017; Sacktor and Fenton, 2018, Pastalkova et al., 2006; Hsieh et al., 2017; Wang et al., 2016)? Remarkably we did not detect differential gene expression-related to these molecules, nor did we detect correlations between any of these molecules and the behavioral measures of avoidance memory expression, whereas we did for *Igf2* as mentioned above. It would however be naive and/or hasty to conclude that despite being unbiased, this transcriptomic screen contradicts the findings of the candidate molecule approaches that have identified a role for these molecules, especially in active place avoidance memory. This highlights one of the shortcomings of using transcriptional profiling to identify molecular mechanisms of functions such as memory. The abundance of mRNA does not relate in a straightforward way to the cellular abundance of protein because the relationship depends on the translational regulation of the particular protein (Vogel and Marcotte, 2012). Indeed, as an example, while PKM*ζ* is strongly regulated by repressing translation at dendrites (Hernandez *et al.*, 2003), there is no evidence that it is transcriptionally regulated, either in late-LTP (Kelly, Crary and Sacktor, 2007) or in active place avoidance conditioning(Hsieh *et al.*, 2017), consistent with the present findings.

While this highlights a confinement of transcriptional profiling, it also draws attention to the necessity of performing combined transcriptional and proteomic profiling in tissue taken from the same subjects. In light of the present findings, the hypothesis that PKM*ζ*’s molecular interactions maintain memory (Sacktor, 2011; Sacktor and Fenton, 2018) predicts finding independence between *Prkcz (*which encodes PKM*ζ*) RNA and measures of memory in an unbiased screen that integrates memory-related measures of behavior with circuit-specific transcriptional profiling as we have done here. The hypothesis also predicts that in the same tissue, there will be concurrent positive relationships between measures of memory and the protein levels of PKM*ζ* and the proteins with which it interacts like KIBRA and NSF (Yao *et al.*, 2008; Vogt-Eisele *et al.*, 2014). There is substantial utility in such analyses that use experimental design and concurrent multi-scale measurements from single subjects in order to integrate information from across levels of biology not only discover unknown and unanticipated mechanisms of complex, poorly defined processes like memory, but to also test the validity of specific mechanistic hypotheses.

## Acknowledgments

This work was initiated in the Neural Systems and Behavior course at Marine Biological Laboratory, to which we are grateful for support and the opportunity to embark on this collaboration. We thank Promega Corporation for generously donating molecular supplies for RNA isolation. We thank Laura Colgin for comments on earlier versions of this manuscript. We thank Becca Young Brim, Caitlin Friesen, Tessa Solomon Lane, Issac Miller-Crews, Taylor Reiter, Amanda Charbonneau, Olga Botvinnik, Lisa Johnson, Erich Schwarz and Titus Brown for critical discussion.

---

**Figure S1.**
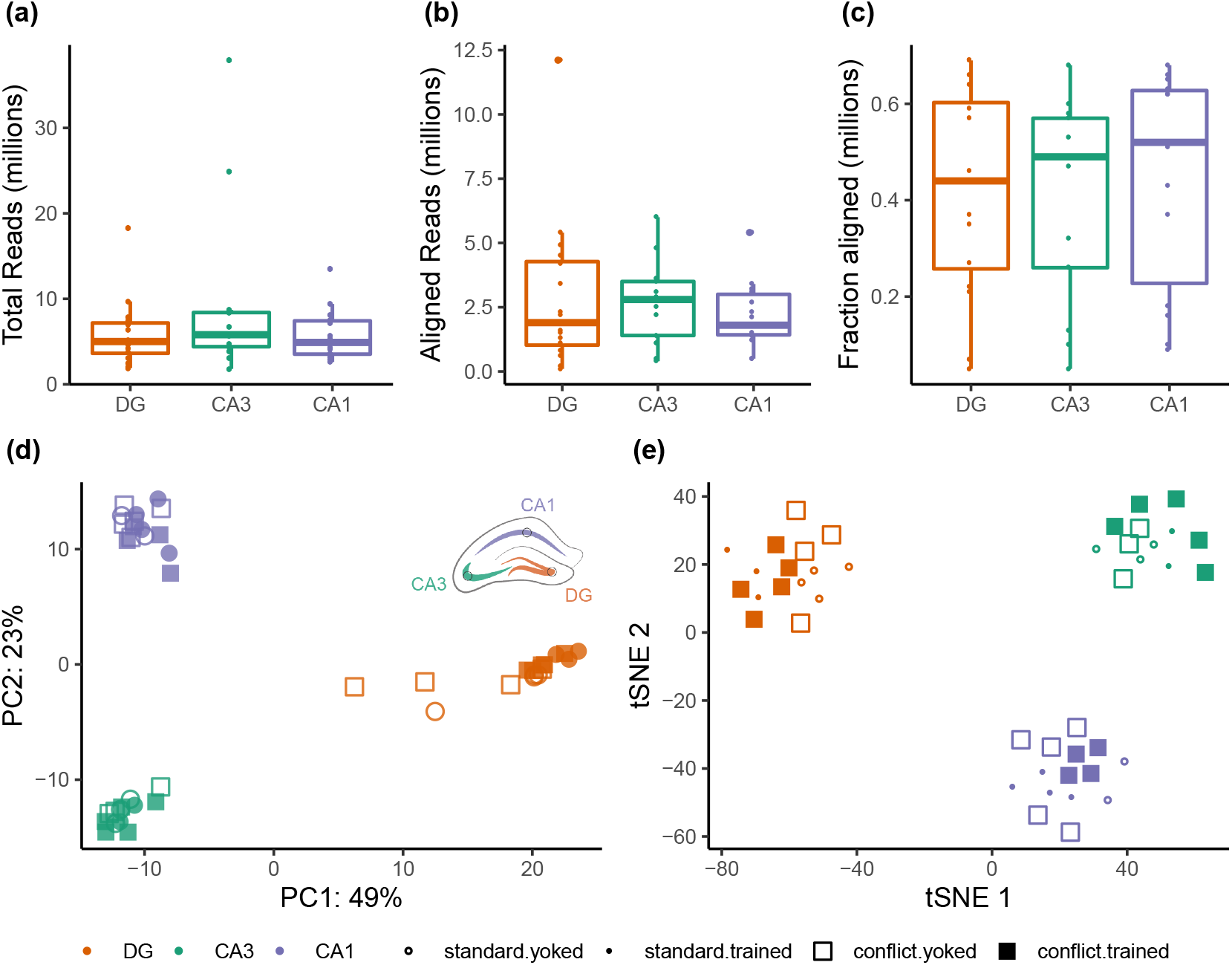
Quality control of RNA-sequencing data. **a)** On average, each sample yielded 6.9 million raw reads. **b)** Of those raw reads, on average 2.6 million mapped to the reference transcriptome. **c)** The alignment efficiency of Kallisto was about 42%. **d)** Principal component analysis (PCA) and **e)** t-Distributed Stochastic Neighbor Embedding (t-SNE) both confirm that all samples can be divided into three clusters based on subfield (DG, CA3, or CA1), as expected, with CA1 and CA3 being more similar to one another than to DG.

**Figure S2.**
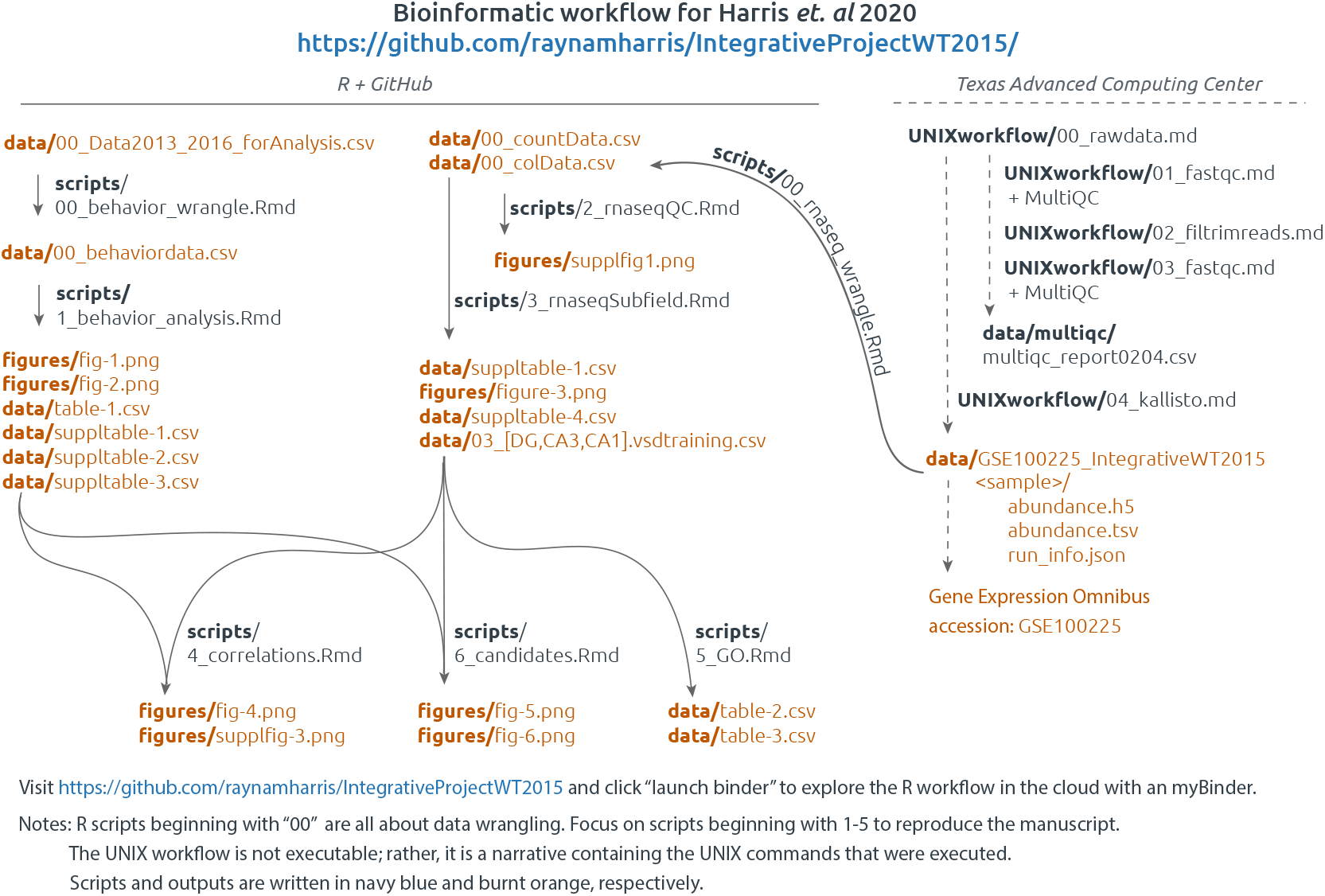
Bioinformatic workflow. This image is a guide for understanding which scripts and data were used to make the figures and tables in this manuscript.

**Figure S3.**
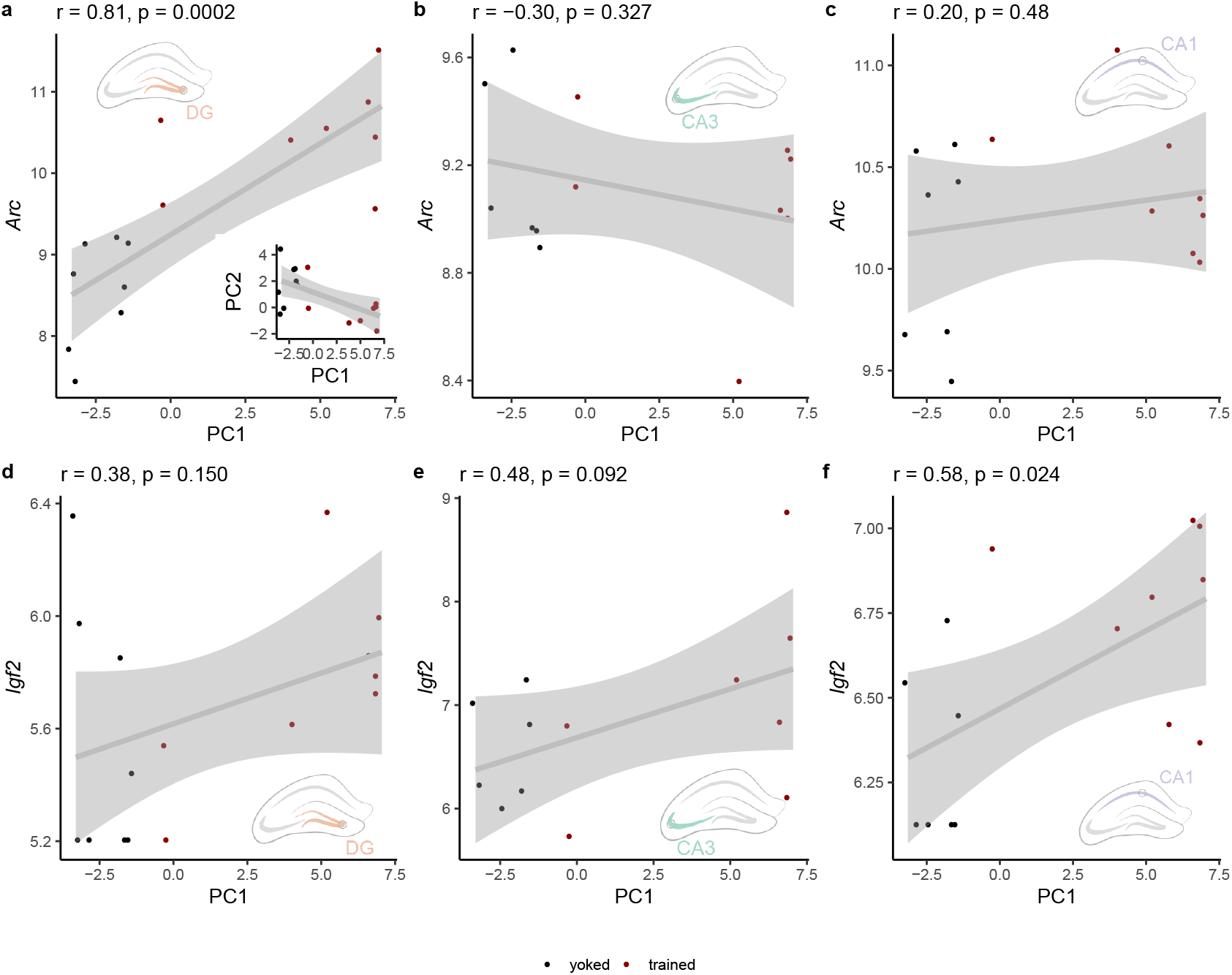
Additional correlations. **a)** *Arc* expression in the DG is significantly correlated with principal components (PC) 1 of the behavioral data. Note, there is a negative correlation between PC1 and PC2 of the behavioral data (inset). **b, c)** *Arc* expression in the CA3 and CA1 is not significantly correlated with PC1 of the behavioral data. **d-f)** Correlations between Igf2 and PC1 are positively in all hippocampal subfields and significant in CA1.

